# The membrane topology problem inherent in ciliogenesis is solved by ciliary recruitment of a phospholipid flippase

**DOI:** 10.1101/2025.01.13.632862

**Authors:** Zhengmao Wang, Ruida He, Qingqing Liu, William J. Snell, Muqing Cao, Junmin Pan

## Abstract

The cilium is a slender cellular projection whose membrane transduces signals essential in development and homeostasis. Although assembly of the axonemal core of the organelle is well-studied, the mechanisms that underlie formation of the ∼250 nm diameter ciliary membrane from oppositely curved 50-100 nm diameter pre-ciliary vesicles remain poorly understood. Here we report that during ciliogenesis in the bi-ciliated alga, *Chlamydomonas reinhardtii*, aminophospholipid flippase 2 (ALA2) is recruited laterally from a reservoir on the plasma membrane onto the nascent ciliary membrane where it transfers conically shaped phosphatidyl ethanolamine (PE) from the outer to the inner leaflet. In cells lacking ALA2, initiation of cilia regeneration after experimentally induced de-ciliation is delayed, PE flipping is impaired, the ciliary membrane becomes distorted, and cilia that do form are shorter than wild type. Our results uncover a membrane topology problem inherent in ciliogenesis and demonstrate that it is resolved by a phospholipid flippase.

**Highlights:**

- Membranes of pre-ciliary vesicles reverse curvature to become the ciliary membrane
- Upon initiation of ciliogenesis, plasma membrane ALA2 transits laterally to cilia
- The ciliary membranes of *ala2* mutants fail to conform to axonemal shape
- ALA2 flips PE to establish ciliary membrane curvature

## INTRODUCTION

The cilium is a slender cellular projection whose membrane mediates multiple sensory functions essential in human development and homeostasis ^1,2^. The canonical, cylindrical shape of a cilium provides a high surface-to-volume ratio for concentrating membrane receptors and maximizing their exposure to the extracellular milieu. Although the formation and function of the microtubule-based core of the cilium, the axoneme, are well-studied, we know little about the mechanisms that control the properties of the lipid bilayer of the ciliary membrane that enable it to enclose the organelle.

During ciliogenesis, the ciliary membrane is formed from the lipid bilayers of pre-ciliary membrane vesicles delivered near the base of the organelle in seamless coordination with assembly of the axoneme ^3,4^. This lipid bilayer provisioning mechanism, however, creates a membrane topology problem: the membrane curvature of the ∼50 - 100 nm diameter pre-ciliary vesicles must be reversed to become the ∼250 nm ciliary membrane. Here, in studies on ciliogenesis in the bi-ciliated green alga, *Chlamydomonas reinhardtii* (hereafter, *Chlamydomonas*), we report a novel and crucial function in ciliogenesis for aminophospholipid flippase, ALA2. At steady state, *ala2* mutant cells either possess short cilia or lack the organelles entirely, and during experimentally induced ciliogenesis, they exhibit a 15-minute delay in initiation of cilia growth compared to wild type (WT) cells. Immunolocalization and immunoblotting show that in WT cells, a full-length, 160 kDa form of ALA2 is present over the entire outer surface of the plasma membrane, with small amounts on the proximal ciliary membrane, whereas a C-terminally truncated 140 kDa form is present in trace amounts on the plasma membrane and enriched in cilia. Both forms become enriched in cilia within the first 15 min of ciliogenesis. The membranes of nascent *ala2* cilia are misshapen, and their outer leaflets are aberrantly enriched in the conically shaped lipid, phosphatidylethanolamine (PE). Our results demonstrate that during ciliogenesis, cells rapidly mobilize ALA2 from the plasma membrane to the ciliary membrane to flip PE and enable the reversal of membrane curvature required to convert pre-ciliary vesicle membranes into ciliary membranes.

## RESULTS

### Ciliogenesis is impaired in *Chlamydomonas* mutants that lack phospholipid flippase, ALA-2, which in WT cells is present on the plasma membrane and on the proximal ciliary membrane

Using CRISPR-Cas9-generated knockout mutants (Figure S1A) for the three phospholipid flippases annotated in the *Chlamydomonas* genome - - ALA1, ALA2, and ALA3 - - we found that only *ala2* cells had a ciliary phenotype. Unlike WT cells, which were nearly 100% ciliated at steady state in the light part of their diurnal LD cycle, ∼30% of the *ala2* mutant cells completely lacked cilia, and the mean length of cilia on the cells that had them was only 70% of that of WT (Figure 1A-1C). Two other, independently derived *ala2* mutants showed similar ciliary phenotypes (Figure S1B and S1C). Notably, cell body morphologies and growth rates of *ala2* cells were like WT - - suggesting a solely ciliary function for ALA2. Experiments to test for redundant functions of the three ALAs showed that cilia of *ala1;ala3* mutants were similar to WT. *ala1;ala2* mutant cells lacked cilia and were present as clusters that had failed to be released from their enclosing mother cell wall after cell division (Figure 1A and 1B), a phenotype characteristic of many aciliated mutants ^5–7^. We failed to obtain *ala2;ala3* mutants. Expression in *ala2-1* mutants of ALA2 with a C-terminal HA tag (ALA2-HA) restored the WT ciliary phenotype (Figure 1A-1C and Figure S1D). Cells with CRISPR/Cas9-generated D410N and E181Q mutations in the ALA2 catalytic core ^8^ (Figure S2A and S2B) exhibited ciliary phenotypes like those of the *ala2-1* null mutant (Figure 1D). Thus, these results show that ALA1 and ALA2 participate in formation or maintenance of cilia, that ALA2 has a dominant function, and that ALA2 phospholipid flippase activity is essential for its function.

**Figure 1.**
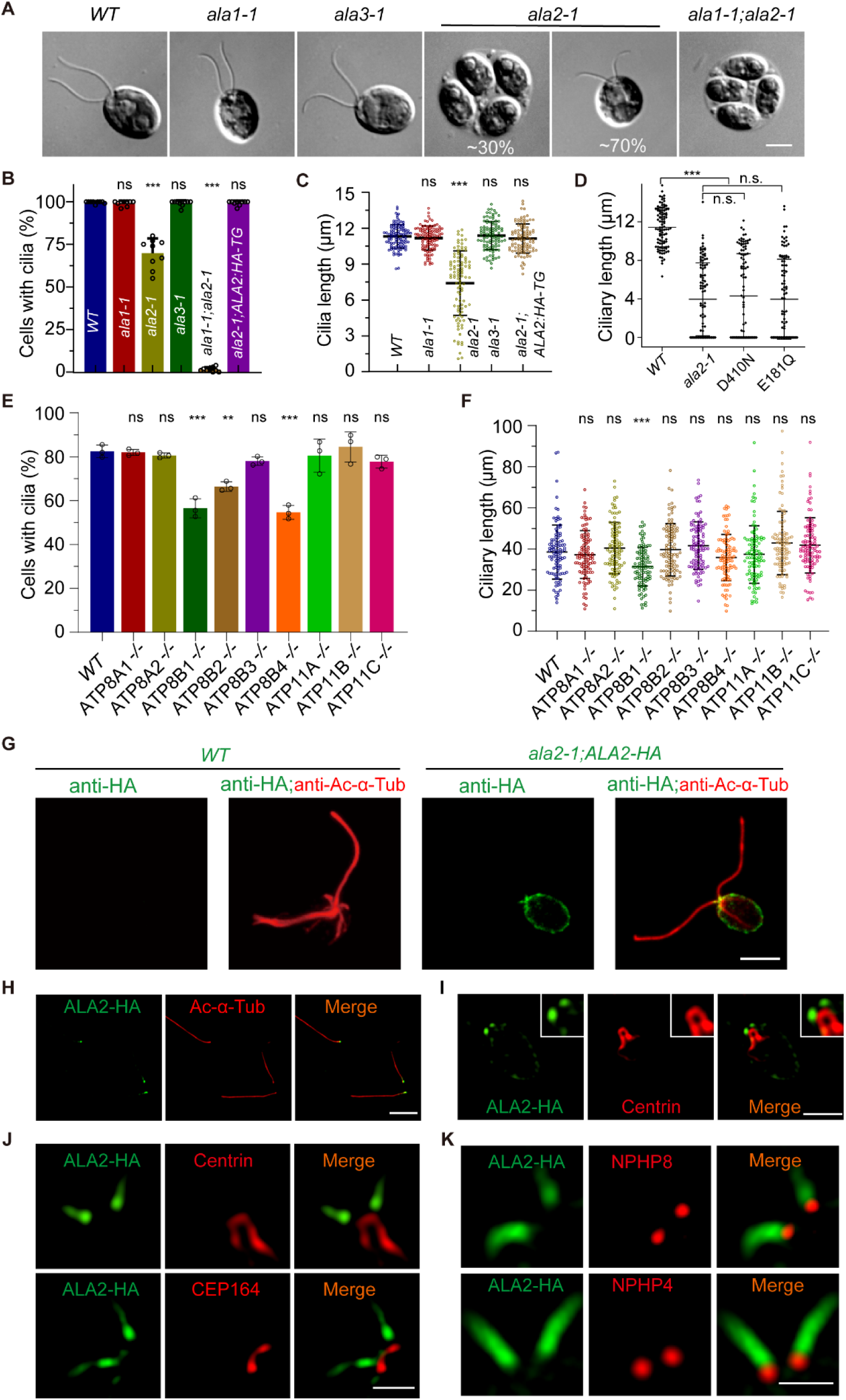
Regulation of ciliogenesis by phospholipid flippases in *Chlamydomonas* and mammalian cells. (A) Representative DIC images of *WT* and mutant *Chlamydomonas* cells as indicated. Bar, 5 μm. (B-C) Ciliation rates (B) and ciliary length (C) from the indicated cells. For ciliation rates, ∼ 500 cells from 9 microscopic fields (40x objective lens) for each strain were analyzed. 100 cells for each strain were analyzed for ciliary length. Error bars, SD. n.s., non-significant; ***, p<0.0001. (D) Defects in ciliary length of the *ala2* catalytic point mutants. Ciliary lengths of 50 cells for each strain from one of two experiments were shown. (E-F) Quantification of ciliation (E) and ciliary lengths (F) of RPE-1 cells mammalian cells. More than 100 cells were counted in each of three experiments. n.s., non-significant; **, p<0.001; ***, p<0.0001. (G) Immunofluorescence images of cells stained with antibodies to HA (green) and acetylated α-tubulin (red). Bar, 5 μm. (H and I) Following deciliation, cilia and cell bodies were immunostained with HA and acetylated α-tubulin antibodies (for cilia) (H) or HA and centrin antibodies (for cell bodies) (I). Bar, 5 μm. (J-K) Co-immunostaining of cells with HA and centrin or CEP164 antibodies (J), and with HA and NPHP8 or NPHP4 antibodies (K). Bar, 1 μm.

To test for a ciliary role of phospholipid flippases in mammalian cells, we disrupted the six genes of the ATP8 group of mammalian flippases in RPE-1 cells that are most closely related to *Chlamydomonas* flippases, and the three genes of the ATP11 group, whose member, *ATP11A*, was present in a primary cilium proteome ^9^ (Figure S3A-3C). Of the nine disruptants, those with disruptions in *ATP8B1*, *B2* and *B4* genes exhibited fewer ciliated cells than WT; ATP8B1 cells also had shorter cilia (Figure 1E and 1F, Figure S3D). Although the ciliary impairments in these single flippase mutants were less severe than those in *Chlamydomonas*, the results suggest that members of this family of phospholipid transfer proteins also function in formation of cilia in mammals.

We used anti-HA immunostaining of the ALA2-HA-rescued *Chlamydomonas ala2-1* cells to determine the cellular location of the tagged protein. In contrast to its apparent function in cilia, nearly all of this C-terminally-tagged ALA2-HA was present on the cell plasma membrane, with only small amounts at the ciliary base (Figure 1G). In samples exposed to a pH shock, which causes the cilia to detach above the transition zone ^10^, the ALA2-HA in the cilia remained at the basal ends (Figure 1H); ALA2-HA at the apical ends of the cell bodies could be seen to be enriched at two puncta (Figure 1I). To determine more precisely the location of the ALA2-HA at the cell body-cilium junction, we used super-resolution microscopy in combination with anti-HA antibodies and antibodies against several transition zone and basal body proteins. These included CEP164, a transition fiber protein ^11^; centrin, a component of the distal striated fiber ^10^, which connects the 2 basal bodies below the transition zone; and two transition zone proteins, NPHP4 and NPHP8. We found that ALA2-HA was present on and above the distal ends of the transition zones (Figure 1J and 1K).

### ALA2 functions early during ciliogenesis, when it rapidly moves laterally from the cell body plasma membrane to become enriched in the proximal ciliary membrane

When *Chlamydomonas* cells are de-ciliated, they draw on a store of precursors from the cell body to initiate immediate growth of new cilia ^12,13^, enabling experimental examination of the relationship between ALA2 and ciliogenesis. In WT cells at 15 min after deciliation, regenerated cilia were nearly 40% of full-length (Figure 2A). In the *ala2-1* mutants, however, cilia were undetectable at 15 min. By 30 min, cilia had appeared on many *ala2* cells, and at 120 min *ala2* cilia had reached about 75% of full-length WT cilia. Consistent with this early ciliogenesis phenotype of the *ala2* mutants, anti-HA staining of ALA2-HA-expressing cells at 15 min after initiation of cilia regeneration showed that the proximal regions of the newly forming cilia had become enriched in ALA2-HA (Figure 2B).

**Figure 2.**
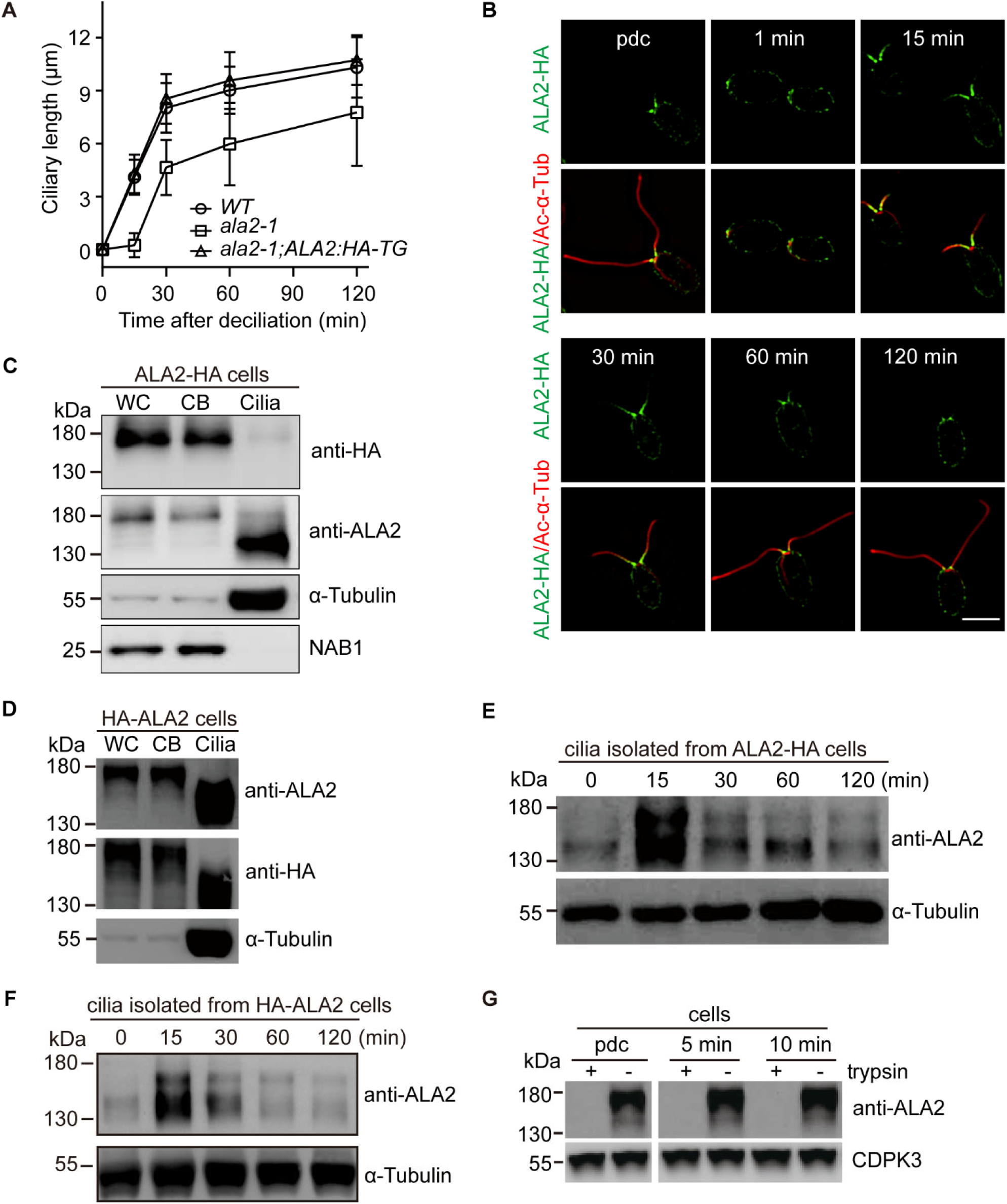
Lateral transport of ALA2 from the plasma membrane to the proximal ciliary membrane during ciliogenesis. (A) ALA2 is required at early steps in ciliogenesis. Ciliary lengths of 50 cells undergoing ciliary regeneration after deciliation (one representative experiment of two). (B) ALA2 becomes enriched at proximal ends of cilia during regeneration. Immunofluorescence images of cells undergoing ciliogenesis at the indicated times after deciliation. Anti-HA (green), anti-α-acetylated tubulin (red). pdc, pre-deciliation; bar, 5 μm. (C-D) Immunoblotting of 10 µg of whole cells (WC), cell bodies (CB), and isolated cilia with the indicated antibodies from cells expressing ALA2-HA (C) and HA-ALA2 (D). Cell body protein loading controls: α-tubulin and NAB1. (E-F) Immunoblotting of 10 µg regenerated cilia isolated from cells expressing ALA2-HA (E) and HA-ALA2 (F) at the indicated times after deciliation. (G) Immunoblotting of trypsin-treated steady-state cells (pdc), and cells 5 and 10 min after deciliation. Loading control: CDPK3, a cytosolic protein kinase.

To confirm these immunolocalization results, we used SDS-PAGE and anti-HA immunoblotting with equal protein amounts (10 μg) of steady-state whole *ALA2-HA* cells and cell body and cilia fractions. ALA2-HA was enriched in cell bodies, and present as a ∼160 kDa molecule, close to the predicted molecular mass of *Chlamydomonas* ALA2 (155,597 kDa) (Figure 2C). These results were consistent with the anti-HA immunofluorescence results of steady-state *ALA2-HA* cells and showed that ALA2-HA was present primarily in the cell body with small amounts in cilia (Figure 2C). Immunoblotting of the same samples with an anti-ALA2 antibody generated against a 161-residue peptide in the cytoplasmic domain (Figure S4), however, showed that cells possessed an additional, 140 kDa, truncated form of ALA2, which was of low abundance in cell bodies, but enriched in cilia (Figure 2C). Anti-ALA2 immunoblots of cells and cell fractions of cells expressing an N-terminally tagged form of ALA2 (HA-ALA2) were similar, confirming that only the C-terminal portion of the protein had been cleaved (Figure 2D).

Anti-ALA2 immunoblotting of cilia isolated from ALA2-HA cells after 15 min of cilia regeneration was consistent with the anti-HA immunofluorescence images of newly regenerating cilia. The 160 kDa form of ALA2 indeed had become enriched in the newly growing organelles. Additionally, though, the 140 kDa form had also become enriched in the cilia (Figure 2E). Similar results were obtained with the cells expressing HA-ALA2 (Figure 2F). Notably, all the ALA2 on live cells before de-ciliation and at 5 and 10 min after initiation of ciliogenesis, was degraded by a brief treatment with trypsin (Figure 2G). Thus, the protein was present on the cell surface in cells at steady state cells and, during ciliogenesis, it remained on the surface during its transit from the plasma membrane to the ciliary membrane. Taken together, these results indicated that cells at steady state possess two populations of ALA2 - - a full-length 160 kDa form localized primarily on the plasma membrane and a truncated, 140 kD form present in trace amounts on the plasma membrane and enriched in cilia. During ciliogenesis, the 160 kDa form undergoes regulated entry into the cilia by lateral movement from the plasma membrane. The 140 kDa ALA2 also appears in cilia early during ciliogenesis, either by lateral movement of pre-existing 140 kDa ALA2 from the plasma membrane, or by cleavage of full length ALA2 during or after transit.

### Transition zone assembly and delivery of vesicles to the base of cilia were unchanged in the *ala2-1* mutant

Although the dual locations of ALA2-HA in the ciliary base and in the transition zone region raised the possibility that the protein also acted in formation of this ciliary membrane barrier, examination of transition zone structure by transmission electron microscopy (TEM) and analysis of the location of the transition zone protein, NPHP8, by immunostaining failed to show changes in *ala2-1* cells (Figure 3A, 3B). We also examined delivery of ciliary vesicles to the ciliary base in WT and *ala2* cell expressing HA-tagged forms of vesicle membrane proteins known to function in ciliogenesis - - syntaxin, a vesicle fusion-mediating t-SNARE, and RAB11, a vesicle trafficking regulator ^14^. Transcriptome analysis showed that the syntaxin transcripts were strongly upregulated in cells regenerating cilia (Figure S5). Immunolocalization studies showed that RAB11-HA and syntaxin-HA were present near the bases of the cilia in WT cells (Figure 3C and 3D). Syntaxin-HA was adjacent to the distal striated fiber protein, centrin, and below the transition fiber protein, CEP164 (Figure 3D), suggesting that ciliary vesicles are delivered just below the transition zone. Most importantly, the locations of the two proteins were similar in WT and *ala2-1* cells (Figure 3C and 3E). Figure 3F illustrates the locations of these and other proteins at the ciliary base. Thus, delivery to the ciliary base of ciliary vesicles whose lipid bilayers are the precursors of the ciliary membrane was independent of ALA2.

**Figure 3.**
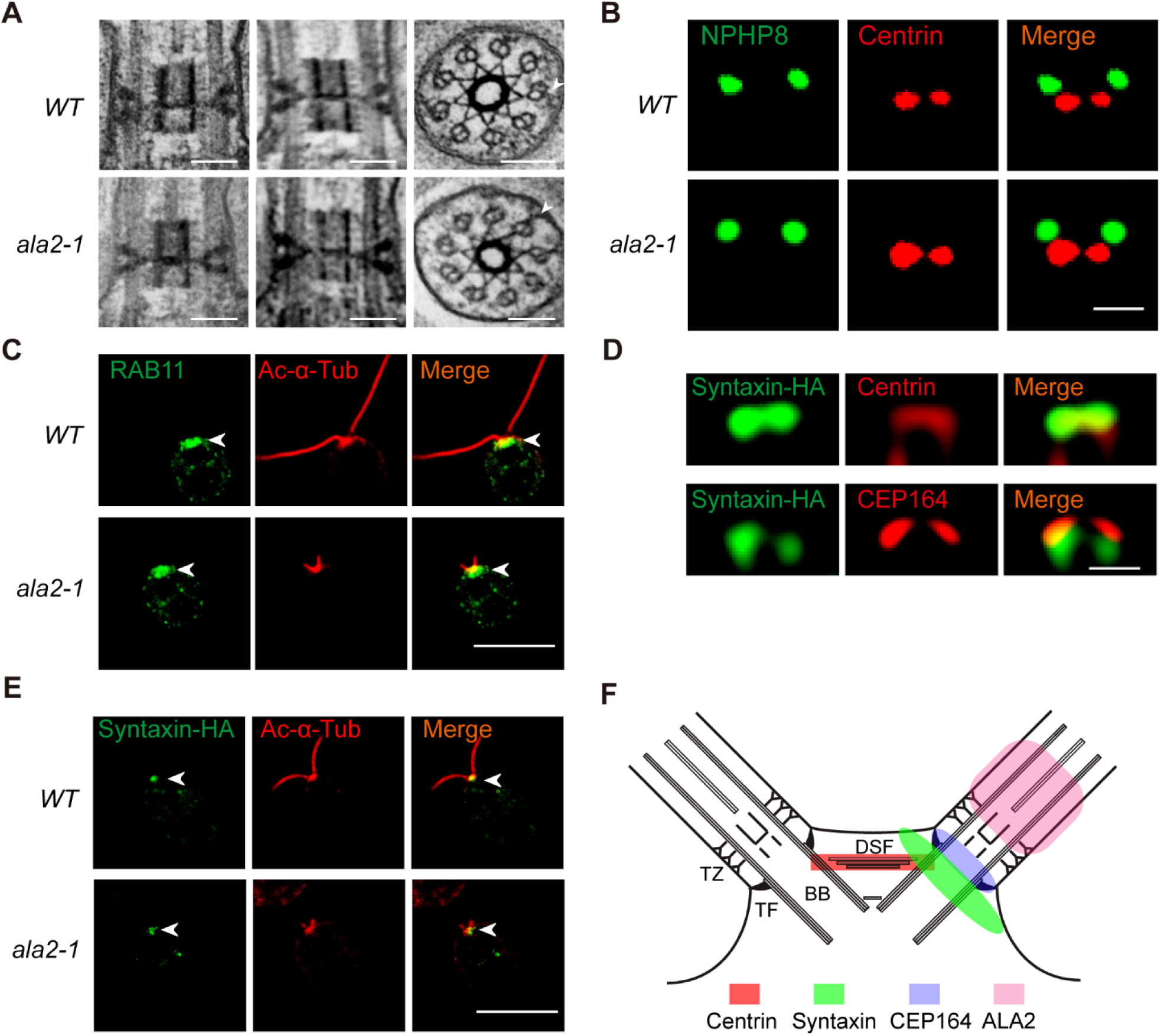
Transition zone assembly and delivery of vesicles to the base of cilia are similar in WT and *ala2* mutants. (A) By TEM, the transition zone of *ala2-1* cells was similar to WT. Arrowheads, Y-shaped links. Bar, 100 nm. (B) Immunostaining shows that the localization of NPHP8 at the transition zone is similar in *ala2-1* and *WT* cells. Bar, 0.5 μm. (C) The delivery of ciliary vesicles (marked by RAB11) to the ciliary base is unaffected in *ala2-1* as shown by immunostaining. Bar, 5 μm. (D-E) HA-tagged syntaxin, a t-SNARE, is positioned adjacent to centrin and proximal to the transition fiber marked by CEP164 (D) and its localization at the ciliary base was not altered in *ala2-1* cells (E) as revealed by immunostaining. Bar, 0.5 μm in (E) and 5 μm in (F). (F) Illustration of protein localizations in structures near the ciliary base. BB, basal body; DSF, distal striated fiber; TF, transition fiber; TZ, transition zone.

### The membranes of *ala2* cilia are misshapen and their outer leaflets are atypically enriched in PE

Examination by transmission electron microscopy (TEM) and scanning EM (SEM) of cells regenerating cilia revealed that, compared to WT cilia, the membranes of *ala2-1* cilia were often constricted at their bases and separated from the axoneme more distally (Figure 4A and 4B). Live cell imaging also showed that newly emerging *ala2-1* cilia became constricted at their proximal ends and detached from the cells (Figure 4C, Supplementary Video 1). Thus, the absence of cilia on many *ala2* cells at steady state likely was a consequence both of a failure to begin to initiate formation of cilia and a failure to retain cilia (Figure 1A and 1B). To determine whether the aciliated phenotype of *ala1;ala2* double mutant (Figure 1A and 1B) cells reflected a more extreme form of cycles of ciliary growth and loss, we examined cultures of *ala1;ala2* strains, whose cells are present in clusters within mother cell walls. Immunostaining with antibodies against acetylated α-tubulin, a marker of ciliary axonemes, showed numerous tubulin-containing structures surrounding the daughter cells (Figure 4D). TEM analysis demonstrated that these structures were intact or partially disrupted axonemes, often without an enclosing ciliary membrane (Figure 4E).

**Figure 4.**
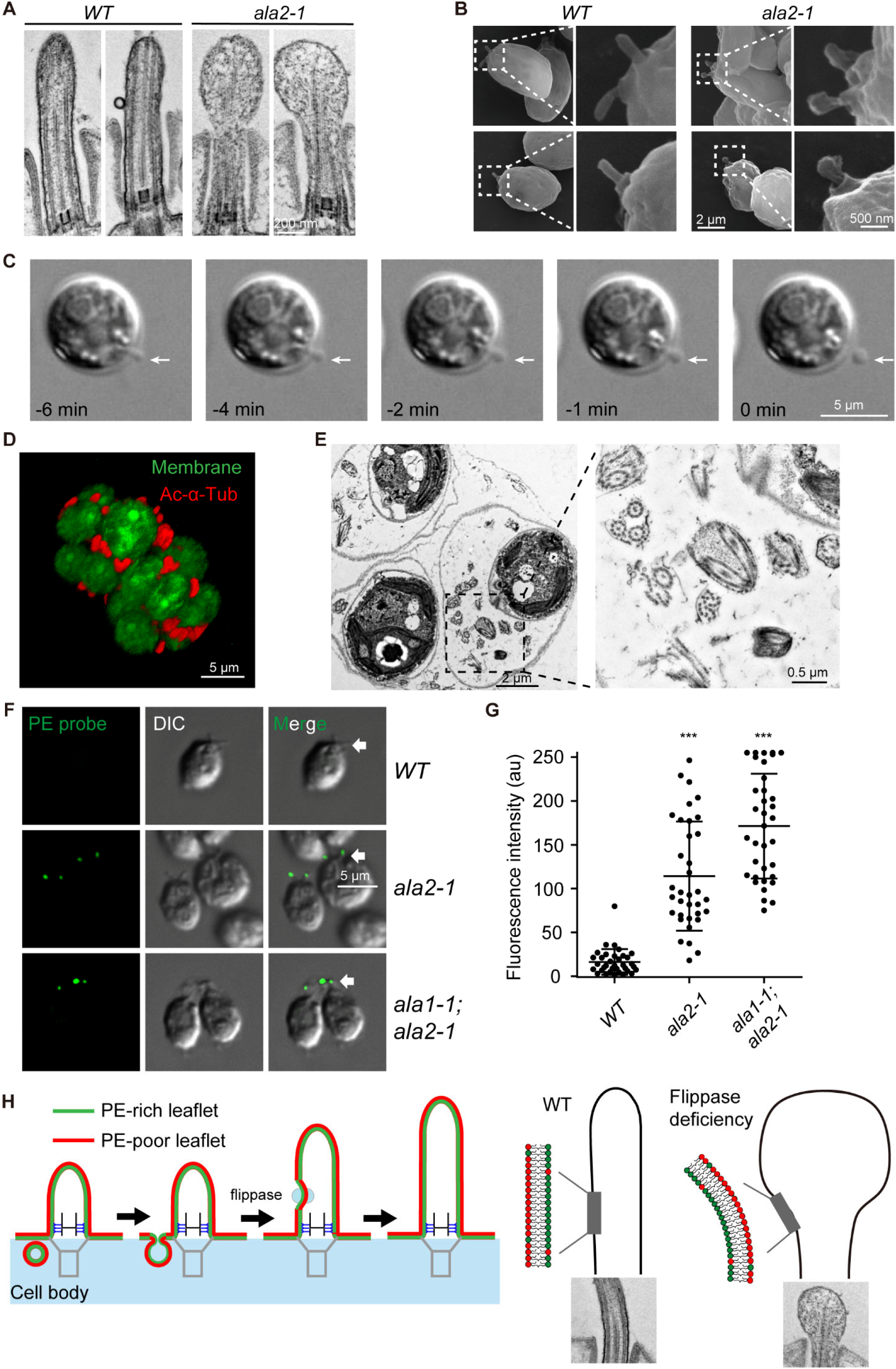
*ala2* nascent cilia are bulbous and the outer leaflets of the ciliary membrane are atypically enriched in PE. (A and B) Representative TEM (A) and SEM images (B) of nascent cilia in *WT* and *ala2-1* cells showing malformed ciliary membrane topology in *ala2-1*. (C) Still images from video of cilia of an *ala2-1* cell forming, swelling, and detaching. Arrows mark cilia. Time 0 denotes the time when cilia detach. (D) The sporangia of *ala1;ala2* mutant were immunostained with anti-α-acetylated tubulin and counterstained with CellMask^TM^ membrane dye. (E) TEM analysis of *ala1;ala2* sporangia showing multiple axonemes surrounding the daughter cells. Boxed area is shown at higher magnification on the right. (F) Representative images showing PE probe labelling of *WT* and *ala2-1* live cells at early stages of ciliogenesis and of steady state *ala1-1;ala2-1* cells. Arrows mark cilia. (G) Quantification of PE signal at the cilia. 30 cilia in each sample were scored. Representative data are shown from one of two experiments. (H) A model for the function of phospholipid flippases in establishing ciliary membrane topology during ciliogenesis. Please see text for details.

Sequence alignments of ALA1 and ALA2 with known phospholipid flippases indicated that both are related to phosphatidyl serine/PE flippases ^15,16^ (Figure S6). PE is a conical phospholipid enriched in the inner leaflets of curved membranes ^17^. As expected, cilia of live WT cells incubated with the PE-specific probe, Cy3-labeled, streptavidin-conjugated biotinylated duramycin ^18^, displayed only trace amounts of PE. On the other hand, the nascent cilia of *ala2-1* and *ala1-1;ala2-1* cells were enriched in PE (Figure 4F and 4G). Thus, the results indicate that phospholipid flippases in the ciliary membrane above the transition zone transfer PE from the outer leaflet to the inner leaflet, thereby establishing the lipid asymmetry and membrane curvature required for proper formation of the ciliary membrane.

## DISCUSSION

Our results show that phospholipid flippases are essential for formation of a stable cilium during ciliogenesis. In the absence of the single flippase, ALA2, *Chlamydomonas* cells fail to generate full-length cilia, and many cells completely lack the organelles. In *ala1:ala2* double mutants, cilia spontaneously detach soon after ciliogenesis begins, and in mammalian cells, flippase disruption is associated with reduced numbers of ciliated cells. The solely ciliary phenotype of the mutants suggests that these ALA flippases function primarily in cilia. Additionally, the normal localization of pre-ciliary vesicle markers at the ciliary base in the *ala2* mutants suggests that vesicle delivery and fusion at the ciliary base are independent of ALA2.

The relatively low amounts of ALA2 in the cilia of steady state cells and the rapid enrichment of ALA2 in cilia in the first 15 min of ciliogenesis (Figure 2) suggest that cells possess a mechanism for coupling the amount of ALA2 in cilia to the rate of delivery of membrane lipids to the organelles. Previous studies reported that cells shed a minimum of 4% of their ciliary membrane every 15 min in the form of ciliary ectosomes ^19^. The WT cells in our studies cilia grew to 40% of full length - - i.e., generating 40% of their ciliary membrane - - in 15 min (Figure 2B). This nearly (40%/4% =) 10-fold increase in rate of delivery of ciliary membrane during ciliogenesis over that being delivered to cilia at steady state is consistent with the large amounts of ALA2 delivered to cilia during this early stage of regeneration.

Our findings that ALA2 on the plasma membrane was the source for the ALA2 that appeared on the ciliary membrane are consistent with previous findings on the source of ciliary membrane proteins in *Chlamydomonas* and mammalian cells and suggest a conserved pathway for mobilizing regulatory membrane proteins into cilia that is distinct from the directed trafficking of Golgi-derived vesicles to the ciliary base ^20,21^ In mammalian cells, the Hedgehog (Hh) pathway effector Smoothened that appears in cilia during pathway activation derives from intracellular stores, but also from lateral movement of pre-existing Smoothened from the plasma membrane ^22,23^. In *Chlamydomonas*, biochemical studies showed that a large, N-terminal fragment (agglutinin) of the *mating type plus* gamete ciliary adhesion protein, SAG1 ^24^, the 350 kDa major ciliary membrane protein, and several unidentified biotinylated proteins ^19^ on the plasma membrane moved onto cilia during ciliogenesis. Additionally, *Chlamydomonas* gametes with full-length cilia at steady state mobilize large amounts of a C-terminal fragment of SAG1 from the plasma membrane onto the ciliary membrane within 5 - 10 min after gamete activation. During mobilization, SAG1 is transiently insensitive to protease treatment, as it is internalized and transported in vesicles along submembranous microtubules to the peri-ciliary membrane followed by exocytosis and re-emergence on the ciliary membrane ^24–27^.

Our finding that the C-terminally truncated, 140 kDa form of ALA2 was enriched in cilia raises the possibility that only this truncated form has catalytic activity. In yeast, the C-terminal segment of phospholipid flippase Drs2p is auto-inhibitory ^28–32^. Based on our findings, it is possible that in addition to regulating the location of ALA2, *Chlamydomonas* also regulates its cleavage, thereby restricting its activity primarily to cilia. It will be interesting to determine whether phospholipid flippases in mammals might be similarly regulated by truncation. We should note that mammalian cells with mutations in inositol polyphosphate-5-phosphatase E (INPP5E) produce misshapen cilia, and that INPPE5 mutations in humans are associated with the ciliopathic MORM syndrome ^33^. Based on these observations and the reports that the truncated yeast Drs2p flippase is stimulated by phosphatidylinositol-4-phosphate (PI4P) ^30–32^, the phosphatidylinositol phosphate products of INPP5E in cilia might also regulate phospholipid flippase activity.

We propose the following model for cilia membrane formation during ciliognesis (Figure 4H). As the organelle extends from the cell, the lipids for its membrane are drawn from lipid bilayers newly delivered near the base by fusion of pre-ciliary vesicles. The inner leaflets of the vesicle membranes become the outer leaflet of the ciliary membrane. Concomitantly, ALA2 that has moved laterally from the plasma membrane to the ciiary membrane, transfers conically shaped PE from the outer leaflet to the inner leaflet to re-establish the lipid asymmetry required for this oppositely curved membrane. Without the flippase, the reversal of membrane topology fails to occur, the ciliary membrane is misshapen and unable to conform to the axoneme, and cilia detach.

Our results that disruption of ATP8B lipid flippases in mammalian cells was also associated with ciliary defects (Figure 1E and 1F) are consistent with this model and could explain phenotypes of flippase mutants in other organisms. The ATP8B protein in *Drosophila* is concentrated in the cilia of olfactory neuron dendrites and its mutation strongly impairs olfactory cell responses to odorants ^34^. Additionally, some human patients with mutations in ATP8B1 showed reduced hearing capacity, and studies in mice suggest that the hearing deficiency could be a consequence of defects in the complex microvilli in hair cells, called stereocilia ^35^. By analogy with our model, generating the high curvature of stereocilia might also depend on a flippase to regulate lipid asymmetry and membrane topology.

### Limitations of the study

Notwithstanding the exciting discovery of the crucial function of phospholipid flippases in the generation and maintenance of ciliary membrane curvature, we note several limitations. The membrane curvature of pre-ciliary vesicles is proposed to be controlled by the asymmetric distribution of phospholipids, with more PE in the inner leaflet of the membrane. Revealing this asymmetry in these intracellular vesicles is technically challenging. Second, we are still uncertain whether the 160 kDa and 140 kD forms of ALA2 are prevented from equilibrating between the plasma membrane and the ciliary membrane by the diffusion barrier at the transition zone or because both are tethered. Such determinations will require use of IFT mutants and more sophisticated live cell imaging. Finally, the site and mechanism of cleavage of the 160 kDa ALA2 to generate the 140 kDa form are unknown. Click chemistry and live cell imaging should help make these determinations. Additionally, the newly available *Chlamydomonas* mutant library should make it possible to screen for protease genes that participate in cleavage.

## SUPPLEMENTAL INFORMATION

Supplemental information can be found online at xxx

## ACKNOWLEDGMENTS

We thank Dr. Liang Wang for kindly providing antibodies. Drs. Liang Ge, Li Yu, Xin Liang for discussions and Lei Liu, Chunlai Chen, Yongxiang Chen and Yanmei Li for help in preparing PE probes. We are grateful to Ying Li, Huizhen Cao and Xiaohui Liu from the Center of Biomedical Analysis (Tsinghua University, China) for technical help. This work was supported by funds from the National Natural Science Foundation of China (31991191, 32370813) to J.P. and (32322021) to M.C.

## AUTHOR CONTRIBUTIONS

Z.W., R.H., Q.L., M.C., W.J.S., and J.P. designed the experiments and analyzed the data. The experiments in *Chlamydomonas* were performed by Z.W. and Q.L. and those in mammalian cells by R.H. and M.C. J.P. and M.C. provided resources. J.P., W.J.S., Z.W., M.C. wrote the manuscript.

## STAR★METHODS

## Key Resources Table

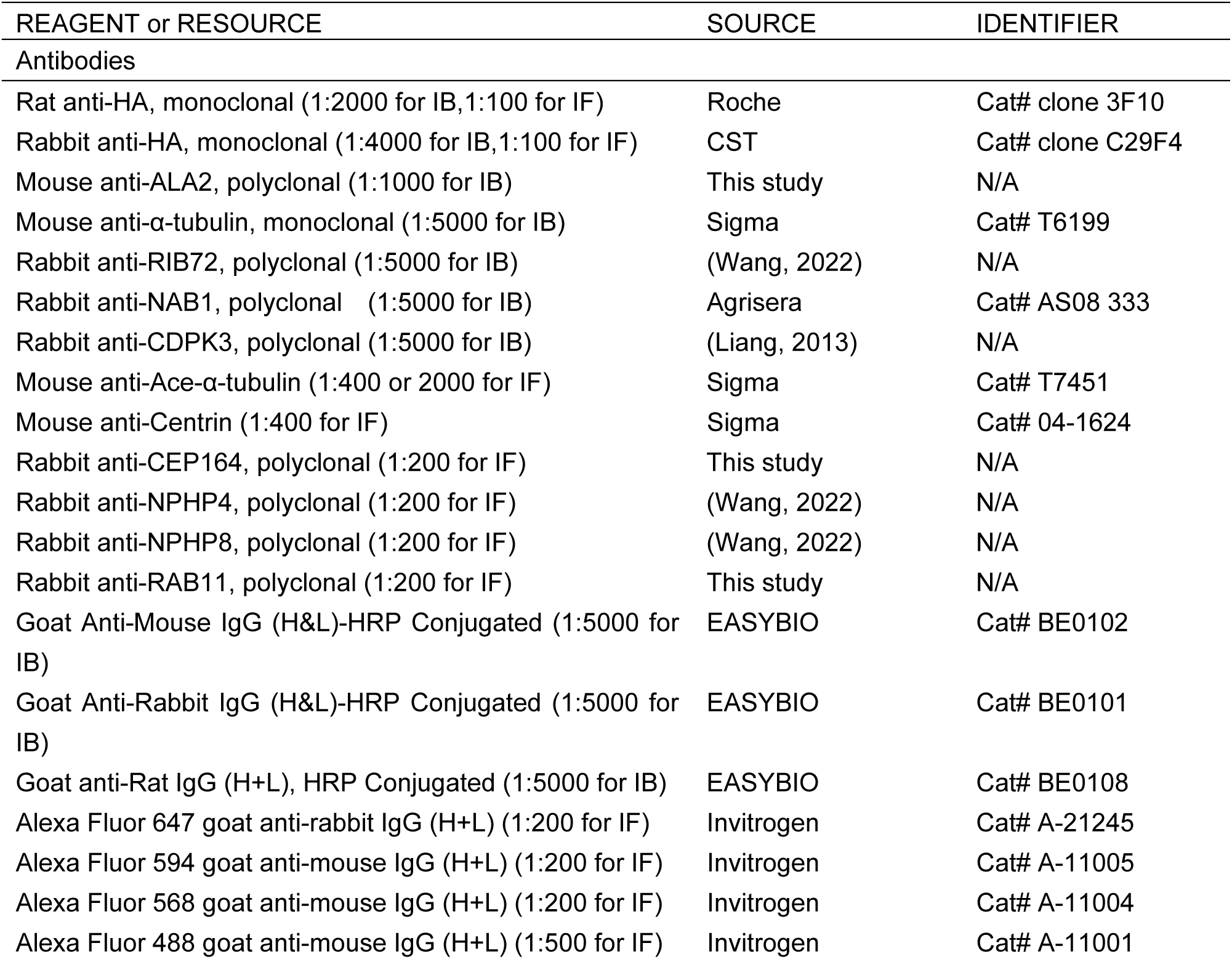
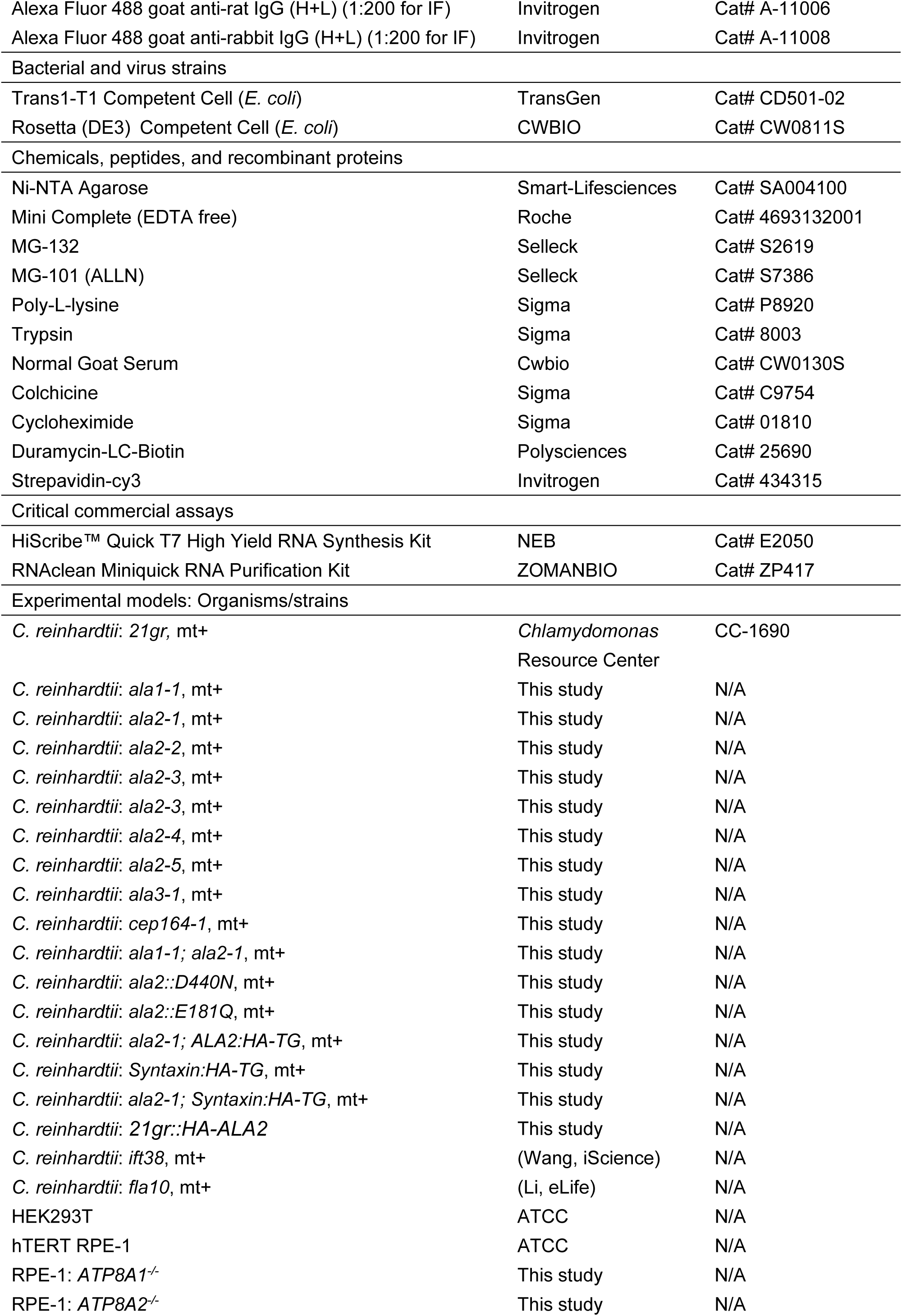
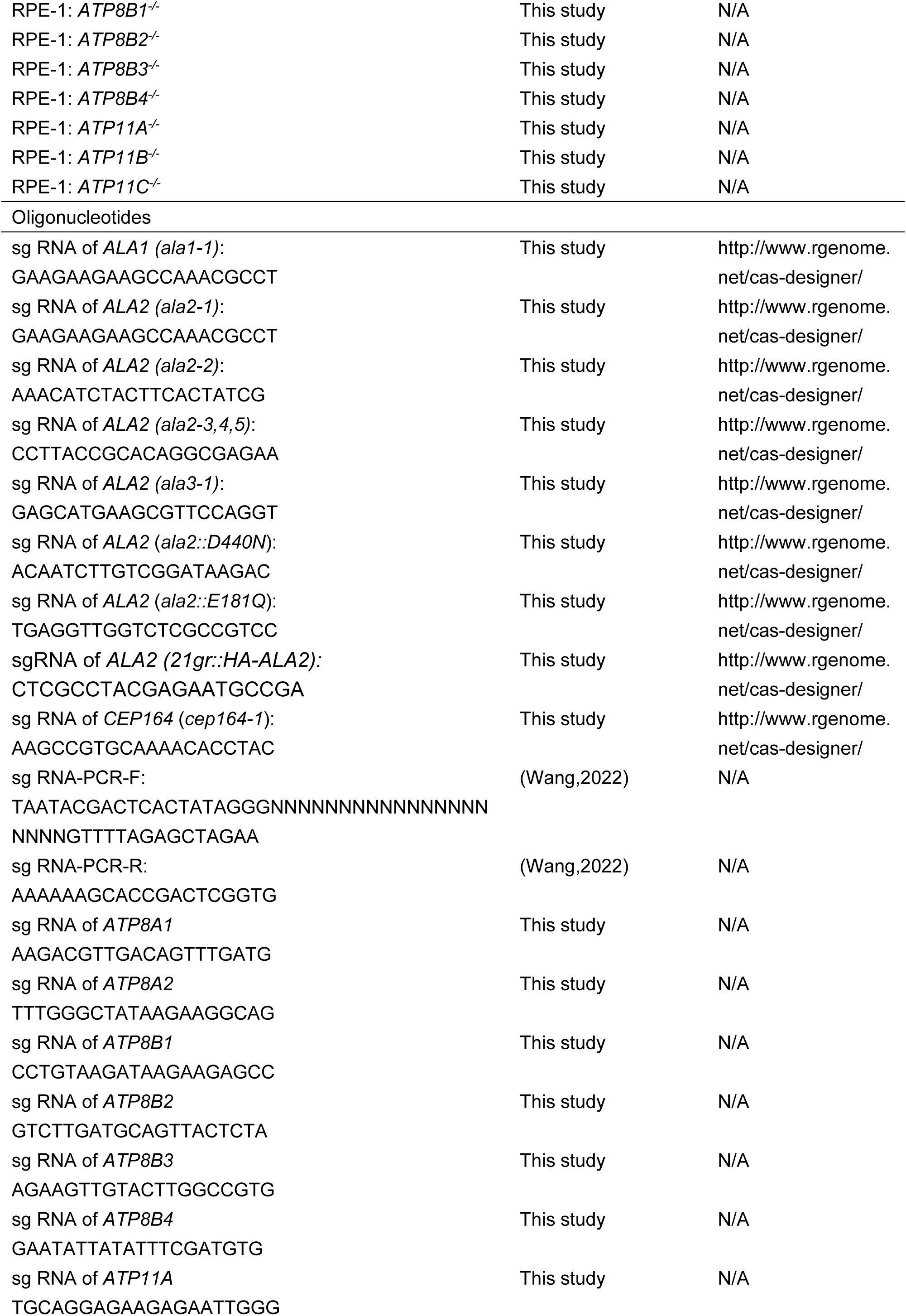
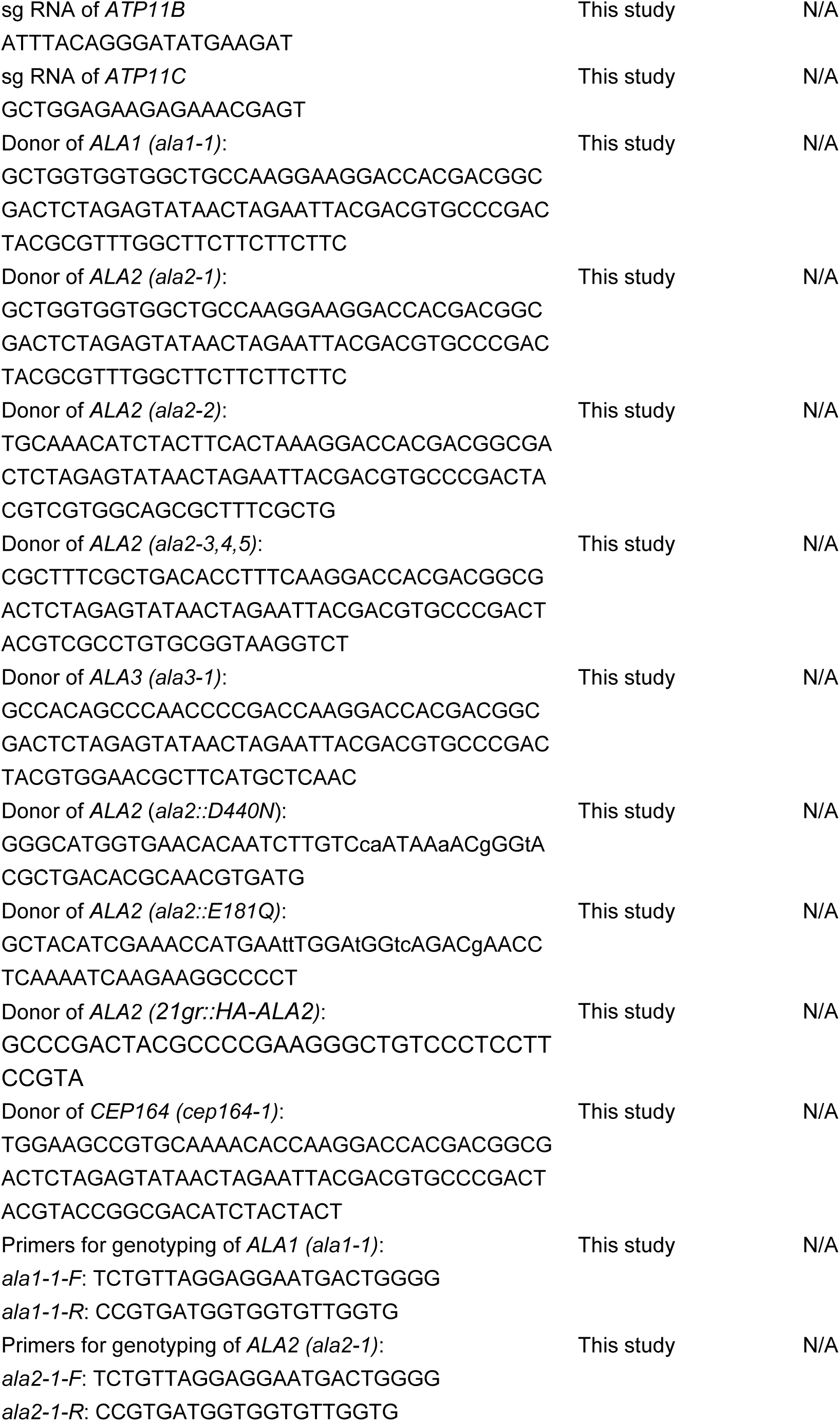
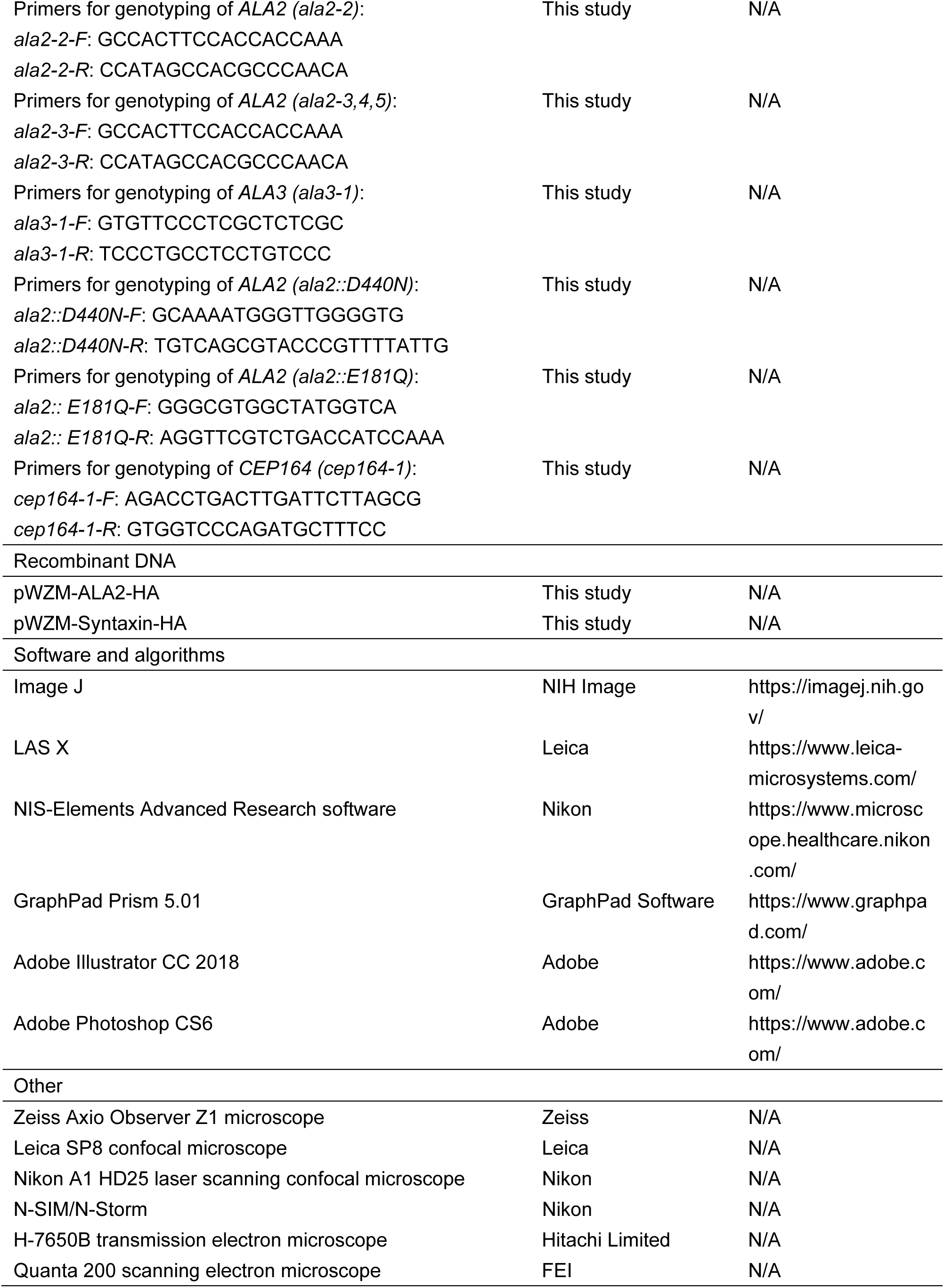

## RESOURCE AVAILABILITY

### Lead contact

Further information and requests for reagents may be directed and will be fulfilled by Junmin Pan, the lead contact.

### Materials availability

*Chlamydomonas reinhardtii* mutants generated in this study will be made available on request, but we may require a payment and/or a completed Materials Transfer Agreement if there is potential for commercial application.

## EXPERIMENTAL MODEL AND STUDY DETAILS

### Strains and cell cultures

*C. reinhardtii WT* strain *21gr* (mt+) (CC-1690), available from the *Chlamydomonas* Genetics Center, was used. All the mutants generated in this study were from this strain background. Cells were grown in liquid minimal (M) medium ^36^ in 200 ml Erlenmeyer flasks with air bubbling on a 14-10 h light-dark cycle at 23 °C. For transformation, cells were grown in liquid tris-acetate-phosphate (TAP) medium ^37^ in constant light. Human hTERT RPE-1 (RPE-1) cells were purchased from ATCC and authenticated. RPE-1 cells were cultured in DMEM/F12 1:1 mixture (Sigma Aldrich, D8437) supplemented with 10% FBS (ExCell Bio, FSP500) and 100 U/ml Penicillin-Streptomycin (GIBCO, 15140122). HEK293T cells purchased from ATCC were cultured in DMEM (Sigma-Aldrich, D6429) supplemented with 10% FBS and 100 U/ml Penicillin-Streptomycin. To induce ciliogenesis in the RPE-1 cells, they were serum-starved by transfer to medium containing 0.5% FBS after having reached approximately 80%–90% confluency. The cells were fixed and stained for analysis after 72 h of starvation.

## METHOD DETAILS

### Genome editing by CRISPR/Cas9 and gene expression in *Chlamydomonas*

Genome editing was performed essentially as previously described ^38^. Briefly, in vitro assembled RNP complex, donor DNA, and a plasmid harboring an expression cassette for paromomycin or hygromycin resistant genes were electroporated together into cells. The colonies growing on selection plates (containing 10 μg/mL paramomycin or 20 μg/mL hygromycin) were screened by PCR followed by sequencing to identify the gene-edited clones.

RNP complex was assembled by mixing 50 μg SpCas9 and 70 μg sgRNA followed by incubation at room temperature for 15 min. For transformation, RNP complex, 1 μg donor DNA and 1 μg of plasmid harboring a selectable marker in a total volume of about 40 μl were mixed with 80 μl cells whose walls had been removed by autolysin ^38,39^. Electroporation was performed in a BTX, ECM630 electroporator (Germany) with the following parameters: 600 V, 50 μF, 300 Ω.

For purification of SpCas9, pPEI-His-SUMO-SpCas9 plasmid was transformed into *E. coli* strain Transetta (DE3) (TransGen Biotech). His-SUMO-SpCas9 bound to Ni-NTA beads was cleaved by Ulp1 overnight at 4 °C followed by purification in a 5-ml HiTrap SP HP Sepharose column (GE Healthcare Life Sciences). SpCas9 eluted with a linear gradient of 100 mM to 1 M KCl was further purified by gel filtration on a Superdex 200 16/300 column (GE Healthcare Life Sciences) in GF buffer (20 mM HEPES, pH 7.5, 150 mM KCl, 1 mM DTT and 10% glycerol). Purified SpCas9 was stored at -80 °C before use.

Single-guide RNAs (sgRNAs) were designed using CRISPR RGEN Tools (http://www.rgenome.net/cas-designer/). sgDNA was generated by PCR based on pX330 plasmid. Synthesis of sgRNA from sgDNA was performed at 37 °C for 5 h using HiScribe^TM^ Quick T7 High Yield RNA Synthesis Kit (NEB, USA, #E2050). For gene knockoutss, donor DNA contained a DNA oligonucleotide with tandem stop codons at the center flanked by 20 bp homology arms from each side of the cleavage site. For point mutations, the donor DNA was prepared to contain the targeted mutations. For rescue of *ala2-1* cells, a plasmid harboring a paromomycin resistance gene and genomic DNA encoding ALA2 with its endogenous promotor and a 3 x HA tag at the 3’ end was introduced by electroporation.

### Transfection, lentivirus infection and generation of mammalian cell lines

Plasmids were transfected into HEK293T cells using linearized polyethyleneimine (PEI) (Polysciences Inc, 24765-2) or Lipocat2000 (Aoqing Biotechnology, AQ11668). The lentiviral knockout vectors were co-transfected with plasmids psPAX2 and pMD2.g into HEK293T cells. The supernatant containing the lentiviral particles was filtered through a 0.45-μm membrane. RPE-1 cells infected with the filtered supernatant were selected with 8 μg/ml of puromycin or 30 μg/ml blasticidin for 2 weeks. Cell colonies were confirmed by Sanger sequencing and Quantitative Real-time PCR (qPCR).

### RNA extraction and Quantitative Real-time PCR

Total RNA was extracted from whole cells using TRIzol^TM^ (Thermo Fisher Scientific, 15596018) following the instructions therein. cDNA was synthesized from 1 μg of total RNA using the SuperScript^TM^ IV First-Strand Synthesis System Kit (Thermo Fisher Scientific, 18091300). Quantitative Real-time PCR (qPCR) was performed using the Universal SYBR Green Fast qPCR Mix (Abclonal, RK21203).

### Analysis of *Chlamydomonas* ciliogenesis and cilia

Cells were deciliated by shearing to allow ciliary assembly as previously described ^13^. Cells were fixed in 1% glutaraldehyde at the indicated times and imaged on a Zeiss microscope (Zeiss Axio Observer Z1) equipped with a QuantEM 512SC camera (Photometrics). Images were exported and processed using ImageJ (NIH), Photoshop and Illustrator (Adobe). Ciliary length was measured using ImageJ (NIH). For each measurement, cilia from at least 50 cells were scored. The microscopy setup described above fitted with a 40x objective was used to capture images of live cells after de-ciliation. Images were acquired at 30 s intervals. Videos were generated using ImageJ (NIH) and played at 2 fps.

Cilia from samples of cells deciliated by a pH shock were purified on sucrose gradients ^13^; suspended in HMDEK buffer (50 mM HEPES, pH 7.2, 5 mM MgCl2, 0.5 mM EDTA, 1 mM DTT, 25 mM KCl) containing protease inhibitors, including protease inhibitor cocktail (mini-complete EDTA-free, Roche, China), 20 mM MG-132 and 65 mM MG-101 (Selleck, China); and frozen in liquid nitrogen before storage at -80 °C.

### Protease sensitivity assay

The protease sensitivity assay was performed essentially as previously described ^27^. Briefly, 1×10^7^ cells were resuspended in culture medium containing 0.05% trypsin at room temperature, harvested after 5 min by centrifugation, washed twice with culture medium containing protease inhibitors and 5% Normal Goat Serum, collected by centrifugation, and flash-frozen in liquid nitrogen.

### Preparation of PE probe and PE cell labeling

To prepare the PE probe, duramycin-biotin (200 μM, Polysciences) was mixed with two volumes of strepavidin-cy3 (1.5 mg/ml, Invitrogen) followed by incubation at room temperature for 1 h. Live cells were incubated with the probe at a 1:200 dilution for 10-30 min at room temperature. After 3 washes with M medium, the cells were fixed with 4% PFA. Cell images were acquired using a Leica TCS SP8 confocal microscope. The fluorescence signals of cilia were calculated using LAX (Leica).

### Primary antibodies, SDS-PAGE and immunoblotting

Details of the antibodies used for immunoblotting or immunofluorescence are described in Key Resource Table. The peptides used for immunizing rabbits to generate polyclonal antibodies were composed of residues 1120-1280 for ALA2, residues 1030-1130 for CEP164, and residues 96-216 for RAB11. All the antibodies were affinity purified (ABclonal and HuaBio, China). Cells and cell fractions were resuspended in HMDEK buffer containing protease inhibitors as described above, boiled in 1x SDS sample buffer for 5 min, followed by analysis of 10 μg portions by SDS-PAGE and immunoblotting.

### Immunofluorescence of *Chlamydomonas* cells

For localization of HA-tagged ALA2, cells were fixed by mixing with an equal volume of 8% paraformaldehyde (PFA) for 5 min. The fixed cells were attached to a PTFE Printed Slide previously coated with 0.01% poly-L-lysine for 10 min. For localization of transition fiber and transition zone proteins, live cells were attached to a PTFE Printed Slide, incubated in -20 °C methanol for 15 min, washed 3 times at room temperature with PBS, blocked with 5% goat serum for 1 h at 37 °C, and incubated at 4 °C overnight with primary antibodies in G-blocking buffer (10 mM PBS, pH 7.4, 1% Gelatin, 0.04% Sodium Azide, 0.5% Goat Serum, 5% Glycerol). After washing three times with PBST buffer (PBS buffer containing 0.5% Tween-20), cells were incubated with the secondary antibodies for 1 h at 37 °C, washed with PBST, mounted with Fluoromount-G (SouthernBiotech), and viewed on a Leica SP8 confocal microscope (Leica) or Nikon A1 HD25 laser scanning confocal microscope using a 100x lens. Images were acquired and processed using LAS (Leica) or NIS-Elements AR Analysis (Nikon) and Photoshop (Adobe). The secondary antibodies used are the following: Alexa Fluor 647 goat anti-rabbit IgG, Alexa Fluor 594 goat anti-mouse IgG, Alexa Fluor 568 goat anti-mouse IgG, Alexa Fluor 488 goat anti-rat IgG and Alexa Fluor 488 goat anti-rabbit IgG from Invitrogen (USA) and all were used at a dilution of 1:200.

For fluorescent quantification of ALA2-HA and acetylated α-tubulin in cilia, the images were analyzed using NIS-Elements AR Analysis (Nikon) and Excel (Microsoft). For structured illumination microscopy (SIM), N-SIM/N-Storm (Nikon) was used. Images were acquired and processed with NIS-Elements AR Analysis (Nikon).

### Immunofluorescence and confocal imaging of mammalian cells

Cells grown on coverslips were fixed with 4% paraformaldehyde (PFA) for 10 min at room temperature, followed by permeabilization in ice-cold methanol for 10 min. After rehydration in PBS for 5 min, and incubated with primary antibodies diluted in blocking buffer (PBS containing 1% bovine serum albumin and 0.1% Triton X-100) for 1 h at room temperature. After washing with PBS, the cells were incubated with secondary antibodies in blocking buffer for 1 h at room temperature. After washing 3 times with PBS, DNA was visualized by 4,6-diamidino-2-phenylindole (DAPI) staining. The coverslips were mounted with FluorSave™ Reagent (Millipore, 345789). Images of the cells were obtained either using an FV3000 Laser Scanning Microscope (Olympus, Japan) equipped with a 40×/NA1.4 oil objective lens (Olympus, Japan). The secondary antibody used was goat anti-mouse IgG Alexa Fluor 488 (1:500; Invitrogen; A-11001) (Luo Genetics in Medicine 2021).

For scoring ciliated cells and ciliary length, three independent experiments were performed. For each assay, at least 100 cells were counted. The average ciliary length was obtained from measuring 100 cilia in each experiment for each strain. The data from one representative experiment are shown.

### Electron microscopy

Cells in M medium were pre-fixed in 1% glutaraldehyde for 15 min at room temperature, harvested by centrifugation, and resuspended in 1% glutaraldehyde, 100 mM sodium arsenate, pH 7.2. After 2 h, samples were incubated with 1% OsO4 in 50 mM sodium cacodylate for 1 h on ice, harvested by centrifugation, and resuspended in freshly prepared 1% uranyl acetate. After overnight incubation at 4°C, the samples were dehydrated, embedded in epon, and thin-sectioned as previously described ^40,41^.

For SEM, cells were first fixed using 1% glutaraldehyde in M medium. Following washes with 100 mM phosphate buffer (PB, pH 7.2) 4 times for 15 min, the samples were post-fixed in 1% OsO4 for 1 h at room temperature. After 4 washes, the cells were dehydrated in a series of ethyl alcohol solutions including 30%, 70%, 80%, 90%, 100%. The cells were placed in 100% tertiary butanol for 15 min, transferred to fresh 100% tertiary butanol for 15 min and freeze-dried in 100% tertiary butanol at -20 °C. The freeze-dried samples were sputter-coated with platinum for 30 s and imaged on a Quanta 200 scanning electron microscope (FEI, Netherlands).

### Bioinformatics analysis

The evolutionary tree of flippases was built using an online tool (http://phylogeny.fr) ^42^. Full length amino acid sequences were used for the analysis. Jalview was used for sequence alignment. InterPro was used for domain analysis.

## QUANTIFICATION AND STATISTICAL ANALYSIS

All the experiments were performed two or more times. The data are presented as means ± SD. Analyses for statistical differences were performed using Student’s t-test with P > 0.05 as non-significant (ns); *, p<0.01; **, p<0.001, ***, p<0.0001.

## Supplemental figures

**Figure S1.**
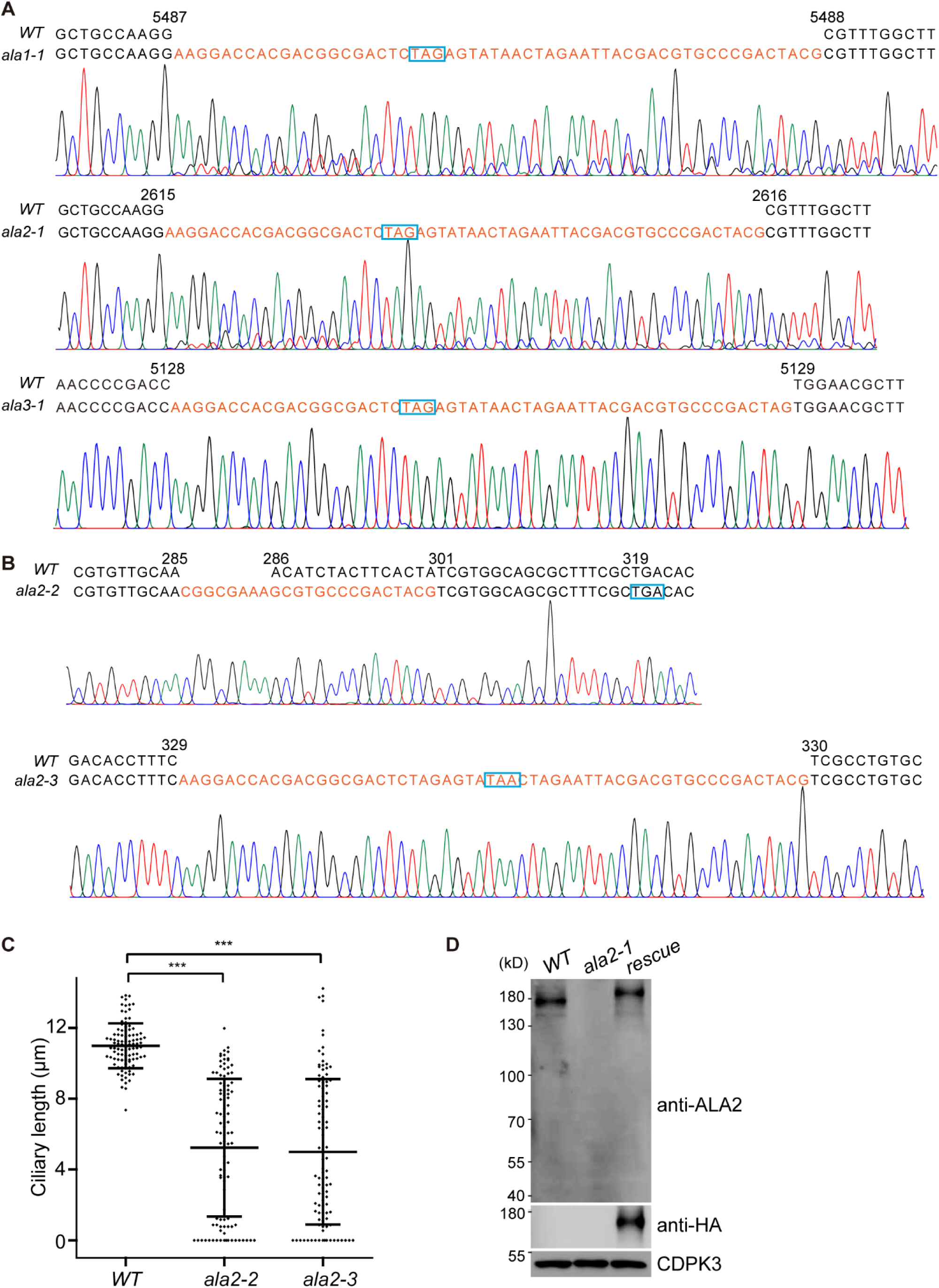
Characterization of *ala* mutants, related to Figure 1. (I) Genome sequences at the edited region of the indicated mutants, which were generated using CRISPR/Cas9. Both the DNA sequence and the sequencing chromatogram are shown. Donor DNA sequences inserted into the genome are highlighted in red, with in-frame stop codons boxed in cyan. (J) Genome sequences at the edited region of two additional mutants of ala2. (K) Ciliary lengths of *ala2-2* and *ala2-3* mutants. Ciliary length from 50 cells for each strain was measured. ***, p<0.0001. (L) Cell lysates from wild-type (WT), *ala2-1,* and rescued cells were subjected to immunoblotting with the indicated antibodies.

**Figure S2.**
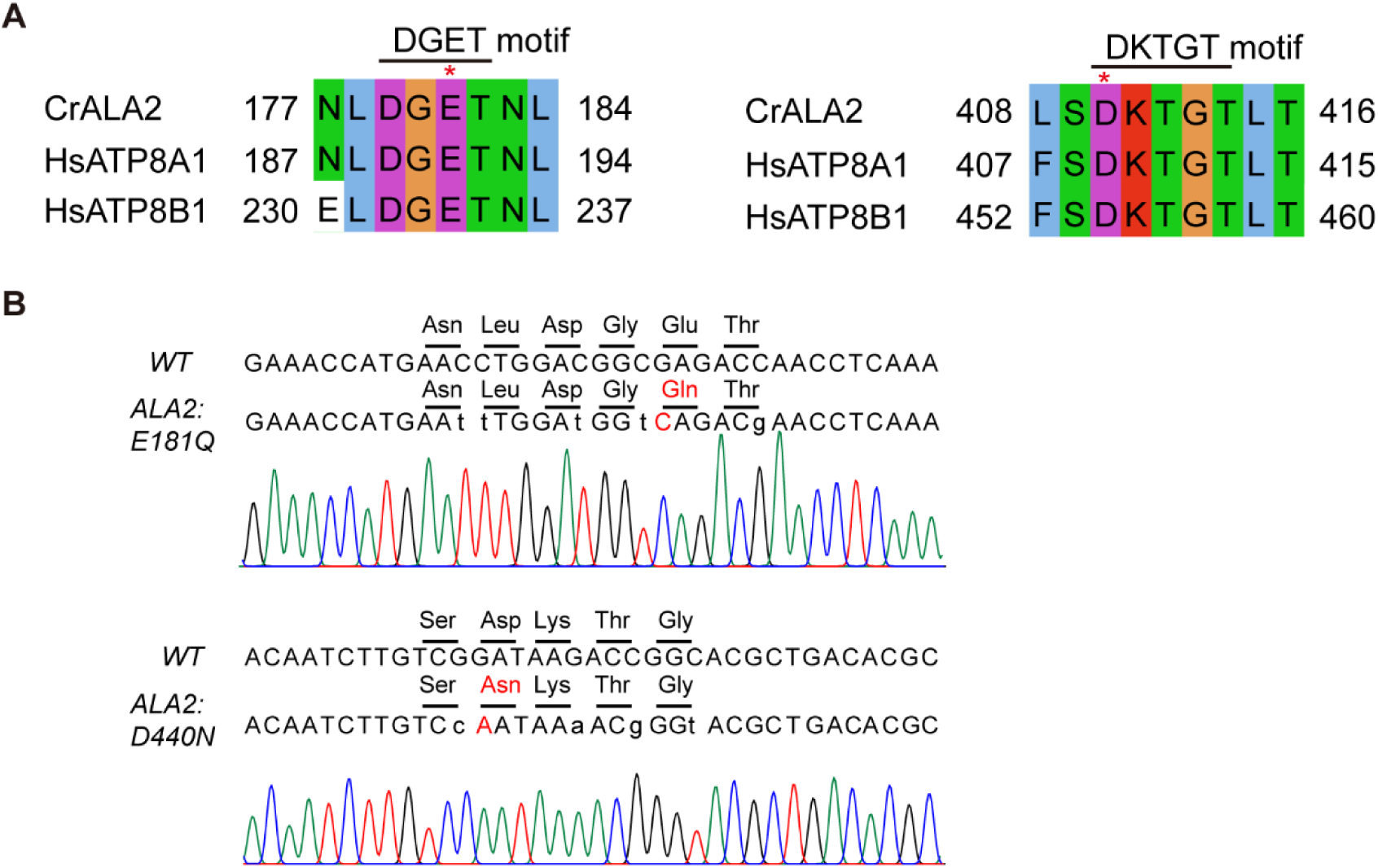
Generation of point mutations at the catalytic domains, related to Figure 1. (A) Sequence alignment of ALA2 with known flippases at the DGET and DKTGT motifs. Stars mark the residues essential for flippase catalytic activity (PMID: 22307598). (B) The catalytic residues of ALA2 were mutated to generate *E181Q* or *D410N* point mutants by gene editing. The residues, nucleotide sequences, and sequencing chromatograms at the edited regions of the mutants are shown.

**Figure S3.**
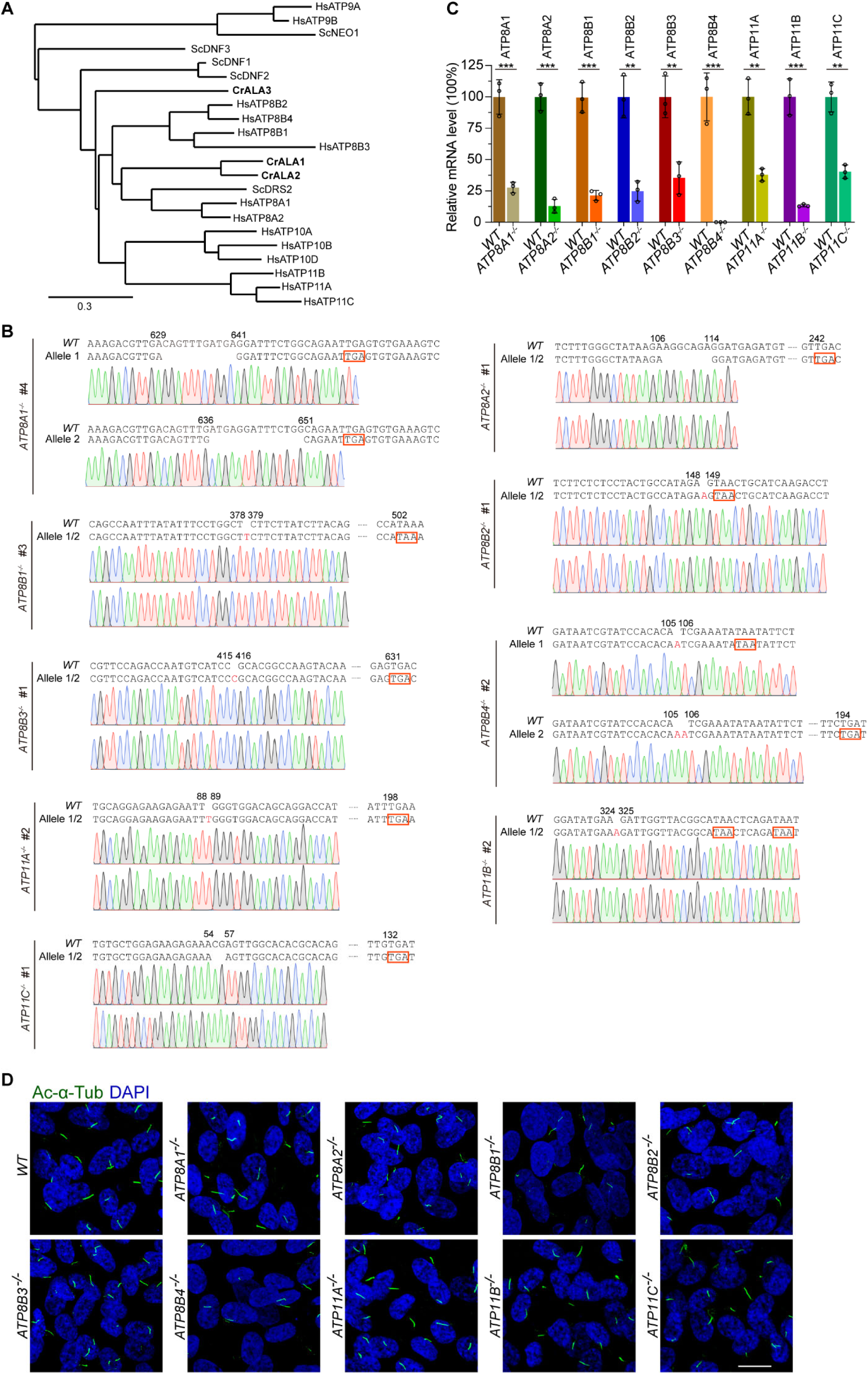
Characterization of phospholipid flippase mutants of RPE-1 cells, related to Figure 1. (A) Phylogenetic analysis of phospholipid flippases from *Chlamydomonas*, human, and yeast, constructed using Phylogeny (www.phylogeny.fr). Hs, *Homo sapiens*; Sc, *Saccharomyces cerevisiae*; Cr, *Chlamydomonas reinhardtii*. Branch lengths indicate evolutionary distance. (B) DNA sequences and sequencing chromatograms of the two alleles in the indicated cell lines. In-frame stop codons are boxed in red. (C) Relative mRNA levels were analyzed by qPCR. **, p<0.001; ***, p<0.0001. (D) Representative images of the flippase knockout cells immunostained for acetylated α-tubulin (Ac-α-Tub, green), and counterstained for DNA (blue). Bar, 20 μm.

**Figure S4.**
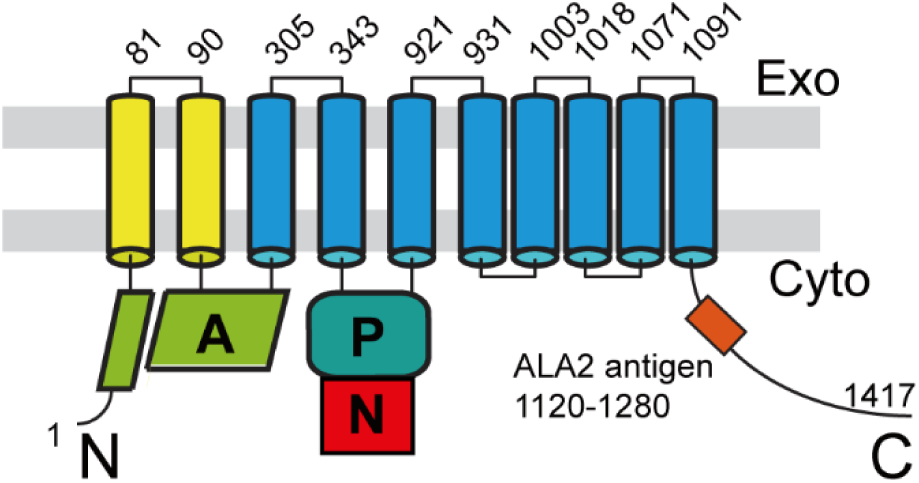
A diagram showing the predicted topology of ALA2, related to Figure 2. The domain structure was predicted using Interpro. The nucleotide-binding (N), phosphorylation (P), and actuator (A) domains are also shown. Exo, exocytoplasmic; Cyto, cytosolic.

**Figure S5.**
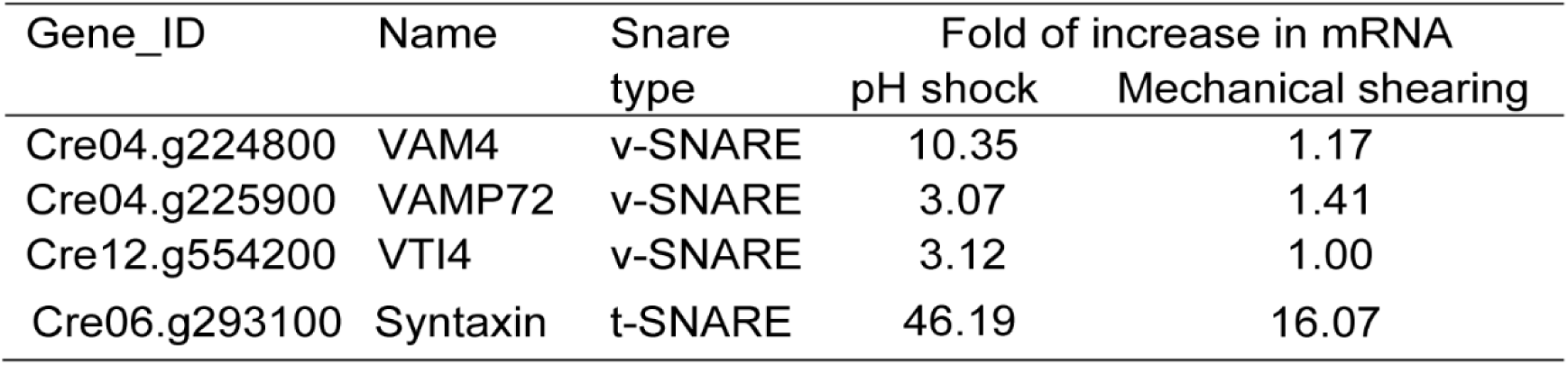
RNA seq data for SNARE genes that are upregulated during ciliogenesis, related to Figure 3. The cells were deciliated either by pH shock or mechanical shearing to allow cilia regeneration. Steady state cells and cells undergoing ciliogenesis at 15 min after deciliation were analyzed by mRNA sequencing. The SNARE genes that showed increased mRNA levels in either of the conditions are shown.

**Figure S6.**
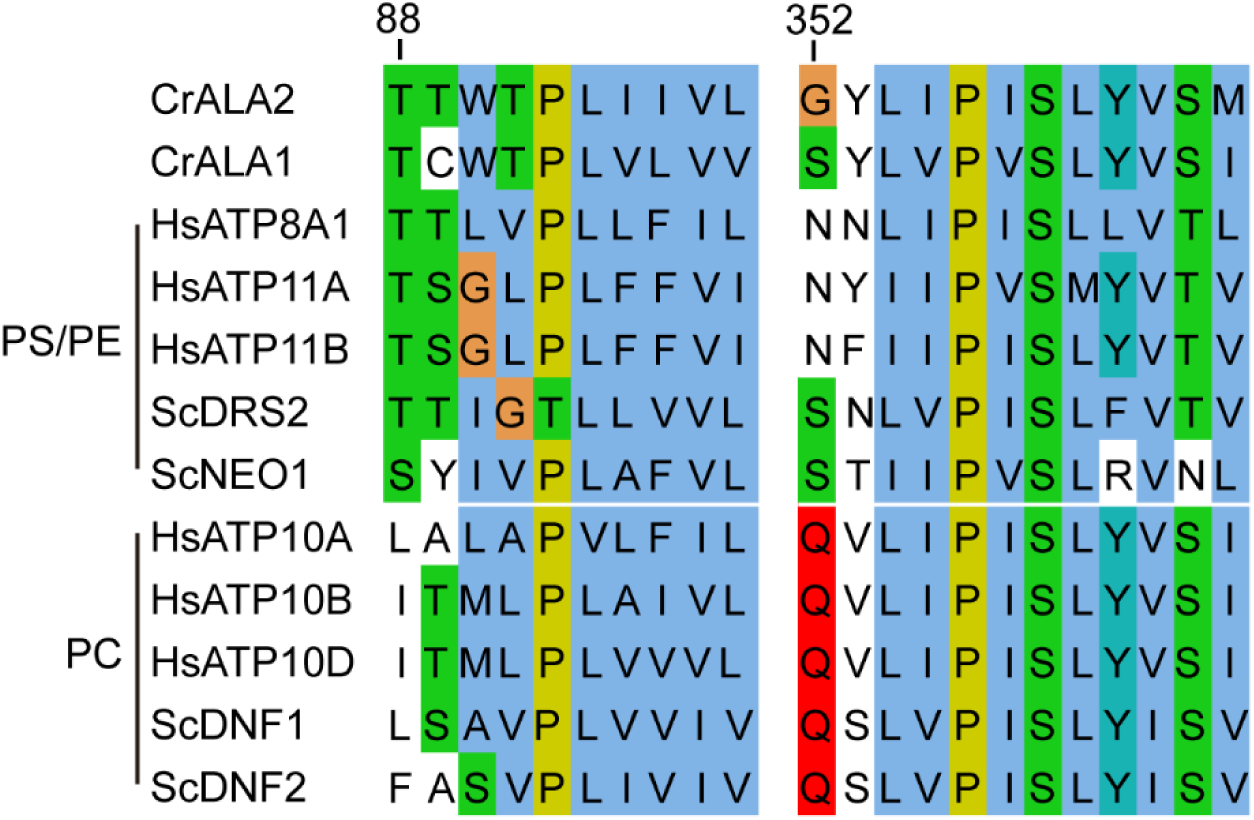
ALA2 is predicted to be a flippase with PS/PE flipping activity, related to Figure 4. Partial amino acid sequence alignment of ALA1 and ALA2 with phospholipid flippases with known flipping PS/PE or PC flipping activities. Please note, threonine (88) of ALA2 is highly conserved among PS/PE flippases while glutamine corresponding to Glycine (352) of ALA2 is conserved among PC flippases.

## REFERENCES

1. Hilgendorf, K.I., Myers, B.R., and Reiter, J.F. (2024). Emerging mechanistic understanding of cilia function in cellular signalling. Nat. Rev. Mol. Cell Biol. 25, 555–573. 10.1038/s41580-023-00698-5.

2. Mill, P., Christensen, S.T., and Pedersen, L.B. (2023). Primary cilia as dynamic and diverse signalling hubs in development and disease. Nat. Rev. Genet. 24, 421–441. 10.1038/s41576-023-00587-9.

3. Shakya, S., and Westlake, C.J. (2021). Recent advances in understanding assembly of the primary cilium membrane. Fac Rev 10, 16. 10.12703/r/10-16.

4. Garcia, G., 3rd, Raleigh, D.R., and Reiter, J.F. (2018). How the Ciliary Membrane Is Organized Inside-Out to Communicate Outside-In. Curr. Biol. 28, R421–R434. 10.1016/j.cub.2018.03.010.

5. Kubo, T., Kaida, S., Abe, J., Saito, T., Fukuzawa, H., and Matsuda, Y. (2009). The Chlamydomonas hatching enzyme, sporangin, is expressed in specific phases of the cell cycle and is localized to the flagella of daughter cells within the sporangial cell wall. Plant Cell Physiol. 50, 572–583. 10.1093/pcp/pcp016.

6. Zhu, X., Liang, Y., Gao, F., and Pan, J. (2017). IFT54 regulates IFT20 stability but is not essential for tubulin transport during ciliogenesis. Cell. Mol. Life Sci. 74, 3425–3437. 10.1007/s00018-017-2525-x.

7. Brown, J.M., Cochran, D.A., Craige, B., Kubo, T., and Witman, G.B. (2015). Assembly of IFT trains at the ciliary base depends on IFT74. Curr. Biol. 25, 1583–1593. 10.1016/j.cub.2015.04.060.

8. Coleman, J.A., Vestergaard, A.L., Molday, R.S., Vilsen, B., and Andersen, J.P. (2012). Critical role of a transmembrane lysine in aminophospholipid transport by mammalian photoreceptor P4-ATPase ATP8A2. Proc Natl Acad Sci U S A 109, 1449–1454. 10.1073/pnas.1108862109.

9. Mick, D.U., Rodrigues, R.B., Leib, R.D., Adams, C.M., Chien, A.S., Gygi, S.P., and Nachury, M.V. (2015). Proteomics of Primary Cilia by Proximity Labeling. Dev. Cell 35, 497–512. 10.1016/j.devcel.2015.10.015.

10. Sanders, M.A., and Salisbury, J.L. (1989). Centrin-mediated microtubule severing during flagellar excision in Chlamydomonas reinhardtii. J. Cell Biol. 108, 1751–1760. 10.1083/jcb.108.5.1751.

11. Tanos, B.E., Yang, H.J., Soni, R., Wang, W.J., Macaluso, F.P., Asara, J.M., and Tsou, M.F. (2013). Centriole distal appendages promote membrane docking, leading to cilia initiation. Genes Dev. 27, 163–168. 10.1101/gad.207043.112.

12. Rosenbaum, J.L., Moulder, J.E., and Ringo, D.L. (1969). Flagellar elongation and shortening in Chlamydomonas. The use of cycloheximide and colchicine to study the synthesis and assembly of flagellar proteins. J. Cell Biol. 41, 600–619.

13. Wu, Q., Gao, K., Zheng, S., Zhu, X., Liang, Y., and Pan, J. (2018). Calmodulin regulates a TRP channel (ADF1) and phospholipase C (PLC) to mediate elevation of cytosolic calcium during acidic stress that induces deflagellation in Chlamydomonas. FASEB J. 32, 3689–3699. 10.1096/fj.201701396RR.

14. Lu, Q., Insinna, C., Ott, C., Stauffer, J., Pintado, P.A., Rahajeng, J., Baxa, U., Walia, V., Cuenca, A., Hwang, Y.S., et al. (2015). Early steps in primary cilium assembly require EHD1/EHD3-dependent ciliary vesicle formation. Nat. Cell Biol. 17, 228–240. 10.1038/ncb3109.

15. Sebastian, T.T., Baldridge, R.D., Xu, P., and Graham, T.R. (2012). Phospholipid flippases: building asymmetric membranes and transport vesicles. Biochim Biophys Acta 1821, 1068–1077. 10.1016/j.bbalip.2011.12.007.

16. Shin, H.W., and Takatsu, H. (2019). Substrates of P4-ATPases: beyond aminophospholipids (phosphatidylserine and phosphatidylethanolamine). FASEB J. 33, 3087–3096. 10.1096/fj.201801873R.

17. Harayama, T., and Riezman, H. (2018). Understanding the diversity of membrane lipid composition. Nat. Rev. Mol. Cell Biol. 19, 281–296. 10.1038/nrm.2017.138.

18. Hullin-Matsuda, F., Makino, A., Murate, M., and Kobayashi, T. (2016). Probing phosphoethanolamine-containing lipids in membranes with duramycin/cinnamycin and aegerolysin proteins. Biochimie 130, 81–90. 10.1016/j.biochi.2016.09.020.

19. Dentler, W. (2013). A role for the membrane in regulating Chlamydomonas flagellar length. PLoS One 8, e53366. 10.1371/journal.pone.0053366.

20. Rohatgi, R., and Snell, W.J. (2010). The ciliary membrane. Curr. Opin. Cell Biol. 22, 541–546.

21. Pazour, G.J., and Bloodgood, R.A. (2008). Targeting proteins to the ciliary membrane. Curr Top Dev Biol 85, 115–149. 10.1016/S0070-2153(08)00805-3.

22. Milenkovic, L., Scott, M.P., and Rohatgi, R. (2009). Lateral transport of Smoothened from the plasma membrane to the membrane of the cilium. J. Cell Biol. 187, 365–374. 10.1083/jcb.200907126.

23. Wang, Y., Zhou, Z., Walsh, C.T., and McMahon, A.P. (2009). Selective translocation of intracellular Smoothened to the primary cilium in response to Hedgehog pathway modulation. Proc Natl Acad Sci U S A 106, 2623–2628. 10.1073/pnas.0812110106.

24. Hunnicutt, G.R., Kosfiszer, M.G., and Snell, W.J. (1990). Cell body and flagellar agglutinins in Chlamydomonas reinhardtii: the cell body plasma membrane is a reservoir for agglutinins whose migration to the flagella is regulated by a functional barrier. J. Cell Biol. 111, 1605–1616. 10.1083/jcb.111.4.1605.

25. Belzile, O., Hernandez-Lara, C.I., Wang, Q., and Snell, W.J. (2013). Regulated membrane protein entry into flagella is facilitated by cytoplasmic microtubules and does not require IFT. Curr. Biol. 23, 1460–1465. 10.1016/j.cub.2013.06.025.

26. Cao, M., Ning, J., Hernandez-Lara, C.I., Belzile, O., Wang, Q., Dutcher, S.K., Liu, Y., and Snell, W.J. (2015). Uni-directional ciliary membrane protein trafficking by a cytoplasmic retrograde IFT motor and ciliary ectosome shedding. Elife 4. 10.7554/eLife.05242.

27. Ranjan, P., Awasthi, M., and Snell, W.J. (2019). Transient Internalization and Microtubule-Dependent Trafficking of a Ciliary Signaling Receptor from the Plasma Membrane to the Cilium. Curr. Biol. 29, 2942–2947 e2942. 10.1016/j.cub.2019.07.022.

28. Zhou, X., Sebastian, T.T., and Graham, T.R. (2013). Auto-inhibition of Drs2p, a yeast phospholipid flippase, by its carboxyl-terminal tail. J. Biol. Chem. 288, 31807–31815. 10.1074/jbc.M113.481986.

29. Natarajan, P., Liu, K., Patil, D.V., Sciorra, V.A., Jackson, C.L., and Graham, T.R. (2009). Regulation of a Golgi flippase by phosphoinositides and an ArfGEF. Nat. Cell Biol. 11, 1421–1426. 10.1038/ncb1989.

30. Timcenko, M., Lyons, J.A., Januliene, D., Ulstrup, J.J., Dieudonne, T., Montigny, C., Ash, M.R., Karlsen, J.L., Boesen, T., Kuhlbrandt, W., et al. (2019). Structure and autoregulation of a P4-ATPase lipid flippase. Nature 571, 366–370. 10.1038/s41586-019-1344-7.

31. Bai, L., Kovach, A., You, Q., Hsu, H.C., Zhao, G., and Li, H. (2019). Autoinhibition and activation mechanisms of the eukaryotic lipid flippase Drs2p-Cdc50p. Nat Commun 10, 4142. 10.1038/s41467-019-12191-9.

32. Herrera, S.A., Justesen, B.H., Dieudonne, T., Montigny, C., Nissen, P., Lenoir, G., and Gunther Pomorski, T. (2023). Direct evidence of lipid transport by the Drs2-Cdc50 flippase upon truncation of its terminal regions. Protein Sci. 33, e4855. 10.1002/pro.4855.

33. Jacoby, M., Cox, J.J., Gayral, S., Hampshire, D.J., Ayub, M., Blockmans, M., Pernot, E., Kisseleva, M.V., Compere, P., Schiffmann, S.N., et al. (2009). INPP5E mutations cause primary cilium signaling defects, ciliary instability and ciliopathies in human and mouse. Nat. Genet. 41, 1027–1031. 10.1038/ng.427.

34. Ha, T.S., Xia, R., Zhang, H., Jin, X., and Smith, D.P. (2014). Lipid flippase modulates olfactory receptor expression and odorant sensitivity in Drosophila. Proc Natl Acad Sci U S A 111, 7831–7836. 10.1073/pnas.1401938111.

35. Stapelbroek, J.M., Peters, T.A., van Beurden, D.H., Curfs, J.H., Joosten, A., Beynon, A.J., van Leeuwen, B.M., van der Velden, L.M., Bull, L., Oude Elferink, R.P., et al. (2009). ATP8B1 is essential for maintaining normal hearing. Proc Natl Acad Sci U S A 106, 9709–9714. 10.1073/pnas.0807919106.

36. Sager, R., and Granick, S. (1954). Nutritional control of sexuality in Chlamydomonas reinhardi. J. Gen. Physiol. 37, 729–742. 10.1085/jgp.37.6.729.

37. Gorman, D.S., and Levine, R.P. (1965). Cytochrome f and plastocyanin: their sequence in the photosynthetic electron transport chain of Chlamydomonas reinhardi. Proc Natl Acad Sci U S A 54, 1665–1669.

38. Wang, J., Zhu, X., Wang, Z., Li, X., Tao, H., and Pan, J. (2022). Assembly and stability of IFT-B complex and its function in BBSome trafficking. iScience 25, 105493. 10.1016/j.isci.2022.105493.

39. Li, X., Zhang, Y., Wen, X., and Pan, J. (2024). Utilizing codon degeneracy in the design of donor DNA for CRISPR/Cas9-mediated gene editing to streamline the screening process for single amino acid mutations. Plant J. 119, 2133–2143. 10.1111/tpj.16903.

40. Craige, B., Tsao, C.C., Diener, D.R., Hou, Y., Lechtreck, K.F., Rosenbaum, J.L., and Witman, G.B. (2010). CEP290 tethers flagellar transition zone microtubules to the membrane and regulates flagellar protein content. J. Cell Biol. 190, 927–940.

41. Meng, D., Cao, M., Oda, T., and Pan, J. (2014). The conserved ciliary protein Bug22 controls planar beating of Chlamydomonas flagella. J. Cell Sci. 127, 281–287.

42. Dereeper, A., Guignon, V., Blanc, G., Audic, S., Buffet, S., Chevenet, F., Dufayard, J.F., Guindon, S., Lefort, V., Lescot, M., et al. (2008). Phylogeny.fr: robust phylogenetic analysis for the non-specialist. Nucleic Acids Res. 36, W465–469. 10.1093/nar/gkn180.

